# A comparison of ThinPrep-PreservCyt against four non-volatile transport media for HPV testing at or near the point of care

**DOI:** 10.1101/2020.03.18.998104

**Authors:** S.G. Badman, A.J. Vallely, C. Pardo, L.P. Mhango, A.M. Cornall, J.M. Kaldor, D. Whiley

## Abstract

**Introduction:** The Xpert HPV Test (Cepheid, Sunnyvale, CA) is used at point-of-care for cervical screening in a number of low-and middle-income countries (LMIC). It is validated for use with ThinPrep-PreservCyt (Hologic, Marlborough, MA) transport medium which has a high methanol content and is therefore classified as a dangerous good for shipping; making cost, transportation and use challenging within LMIC. Therefore, we compared the performance of ThinPrep against four non-volatile commercially available media for HPV point-of-care testing.

**Methods:** Ten-fold serial dilutions were prepared using three HPV cell lines each positive for 16, 18 or 31 and with each suspended in five different media types. The media types consisted of Phosphate Buffered Saline (Thermo Fischer Scientific, Waltham, MA, USA), Sigma Virocult (Medical Wire & Equipment, Wiltshire, England), MSwab (Copan, Brescia BS, Italy) Xpert Transport Media (Cepheid, Sunnyvale, USA) and ThinPrep-PreservCyt (Hologic Inc., Marlborough, MA).

**Results:** A total of 105 HPV Xpert tests were conducted in a laboratory setting, with 7 ten-fold dilutions of each of the 3 HPV genotypes tested in all 5 media types. The lowest HPV ten-fold dilution detected for any media, or cell line was the fifth dilution. Mswab was the only media to detect HPV to the 5^th^ dilution across all three cell types.

**Discussion:** Mswab transport media may be a suitable alternative to ThinPrep for Xpert HPV point of care testing and increase HPV detection. A field-based, head to head comparison of both media types using the Xpert HPV assay is warranted to confirm these laboratory-based findings.

## Introduction

An estimated 570,000 cases of cervical cancer and 311,000 cancer-related deaths occur every year worldwide.(1) More than 85% of this burden is borne by women in low-and middle-income countries (LMICs) where HPV vaccination, cervical screening, and effective treatment are largely unavailable.(1) In high-income settings, the cytology-based Papanicolaou (Pap) test transformed cervical cancer control and led to major reductions in cancer incidence and mortality.(2) The test is, however, technically difficult and requires extensive infrastructure for quality control and referral.

In the last decade, the effectiveness of HPV nucleic acid testing as a screening strategy for the detection of cervical pre-cancer and cancer has been demonstrated in randomised trials and prospective studies,(3-4) leading the WHO to recommend it for primary screening.(5) In high-income countries, HPV testing is now replacing cytology as the preferred first step in screening. For LMICs, however, it has not been considered feasible due to the cost of assays and laboratory equipment, and the lack of laboratory and clinical capacity required for testing, follow-up and treatment. A further significant breakthrough in the last 5 years has been the development of HPV test platforms with the potential for use at point-of-care (POC) in routine clinical settings in LMICs.

The Xpert HPV Test (GeneXpert; Cepheid, Sunnyvale, CA) is a cartridge-based, non-batch test that has been extensively evaluated against other laboratory-based HPV molecular assays and found to have high sensitivity and specificity for the detection of underlying high-grade disease using both clinician and self-collected samples.(6-8) Xpert HPV has been validated for use with ThinPrep-PreservCyt Solution (Hologic Inc., Marlborough, MA): using clinical specimens (e.g. cytobrush, swab) collected in a 20mL vial of ThinPrep solution to suspend cellular material; a 1 mL aliquot is placed in the Xpert Test cartridge prior to testing on the GeneXpert instrument.(9) This test is approved for primary cervical screening in Australia. The composition and unit cost in some countries of ThinPrep are however important constraints to the introduction and scale-up of POC Xpert HPV testing for cervical screening in LMIC settings. Each vial contains up to 60% methanol making it flammable and toxic.(10)

Organic compounds such as ethanol are also known PCR inhibitors, and the concentration of the compound may influence its inhibitory affect.(11) Other studies have suggested DNA may degrade over time when stored in this media.(12-14)

The International Air Transportation Association (IATA) categorises this product as a UN1993 Dangerous Good (DG).(10) This classification incurs additional costs (above the initial purchase price), due to the requirement for specialised handling and packaging before air transportation.(15) These additional cost and logistics issues are considered major challenges to establishing and maintaining the necessary supply chains required to scale-up HPV-based cervical screening services in LMICs.

In this paper, we present findings from a laboratory study of the performance of Xpert HPV for the detection of known HPV cell lines (16, 18 and 31) suspended in ThinPrep and four commercially available and non-volatile PCR transport media.

## Methods

### HPV cell types

HPV-positive cell line material was obtained from the following sources. CaSki [American Type Culture Collection (ATCC) CRL-1550], containing the HPV16 genome {Pattillo, 1977 #852}, were cultured in RPMI-1640 medium with 10% foetal bovine serum (FBS). HeLa (ATCC CCL-2), containing the HPV18 genome {Scherer, 1953 #853}, were cultured in Eagle’s Minimum Essential Medium with 10% FBS. CIN-612 cell line, containing the genome of HPV31 subtype b {Bedell, 1991 #854}, were generously provided by Laimonis A. Laimins, Feinberg School of Medicine, Northwestern University, Chicago IL (USA). All cell lines were suspended in <1mL of ThinPrep-PreservCyt medium (Hologic, Inc., Marlborough MA, USA).

### Human DNA

The Xpert HPV Test contains sample adequacy control (SAC) reagents to detect the presence of a single copy human gene and determine if specimens contain adequate cellular material for testing and minimising false-negative results. Human genomic DNA purchased from Bioline Australia (Cat No: BIO-35025) was spiked at a static concentration to each of the HPV serial dilution samples to provide valid SAC results.

### Media selection

Four commercially available non-volatile transport media suitable for PCR were chosen to compare against ThinPrep-PreservCyt (Hologic Inc., Marlborough, MA): Phosphate Buffered Saline (PBS) (Thermo Fischer Scientific, Waltham, MA, USA), Sigma Virocult (VTM) (Medical Wire & Equipment, Wiltshire, England), MSwab (Copan, Brescia BS, Italy) and Xpert Transport Media (TM) (Cepheid, Sunnyvale, USA).

### Laboratory procedures

For each media type, a serial dilution was prepared using HPV cell lines 16, 18 and 31, respectively. Seven 2mL screw-top microtubes (Biologix®, Shandong, China) were each filled with 1.4mL of each media type, and 10μL of the pre-prepared purified human DNA was added to each of those tubes. A 15 μL volume of the relevant HPV cell line was added to the first tube in each series, and the contents mixed thoroughly by vortex. From that first vial, 140 μL of media mix was transferred to the second vial, which was also vortexed. The process was repeated sequentially for the remaining 5 tubes, creating seven 10-fold dilutions of HPV cells. This process was repeated for each remaining cell line and each of the 5 media types. A 1mL aliquot from each tube was then transferred by disposable pipette to individual Xpert HPV test cartridges and tested as per manufacturer’s instructions.

## Results

A total of 105 Xpert HPV tests were conducted in a laboratory setting, with 7 ten-fold dilutions of each of the 3 viruses tested in all 5 media types (Table 1). The SAC was detected in all samples. The lowest HPV ten-fold dilution detected for any media or HPV type was the fifth dilution, with dilutions 6 and 7 providing negative results for all cell lines and all media types.

Overall the Mswab was the only media that provided detection of HPV to the 5^th^ dilution for all cell lines. The remaining media failed to detect the 5^th^ or in some cases, lower dilutions for one or more cell lines.

There were some anomalous results observed for the HPV 31 dilution series; for Viral TM, HPV was not detected (ND) in step 4 but was detected again in step 5 (Ct 32.3); likewise, for Xpert TM media, HPV was not detected in dilution 2 and was detected again in dilution 3 (Ct 31.8). These variations were not further investigated.

## Discussion

Overall, our results suggest the Xpert HPV test was consistently more sensitive in the detection of HPV cell lines 16, 18 and 31 when suspended in Mswab media. Thus, these preliminary data suggest Mswab media may be a viable alternative to ThinPrep when testing for HPV at or near the POC. A reduction in media vial volume (20mL for Thinprep versus 2.3 mL for Mswab) may also increase sample cell density and improve HPV detection.

One limitation of the study is the use of cell line products instead of clinical samples. The emulation of clinical samples in each dilution step does not account for the presence of natural inhibitors found in clinical specimens. A second limitation is that the HPV tests were not run in duplicate or triplicate, and we do not know how reproducible the results were.

A large-scale head to head comparison of both media types using the Xpert HPV assay is warranted to confirm these laboratory-based findings and determine if this product is suitable for HPV POC testing and scale-up.

## Access to data

All data generated in this evaluation has been declared in this manuscript.

## Contribution

SB designed this evaluation in conjunction with DW. Laboratory experimentation was led by SB in conjunction with CP, LM and AC. Editorial refinement was conducted by AV and JK.

---

**Table S1.** Comparison of ThinPrep-PreservCyt against four commercial non-volatile PCR media types.

| Dilution | Viral TM |  | PBS |  | Xpert TM |  | Thinprep TM |  | Mswab TM |  |
| --- | --- | --- | --- | --- | --- | --- | --- | --- | --- | --- |
| HPV 16 | SAC<br>(Ct value) | HPV<br>(Ct value) | SAC<br>(Ct value) | HPV<br>(Ct value) | SAC<br>(Ct value) | HPV<br>(Ct value) | SAC<br>(Ct value) | HPV<br>(Ct value) | SAC<br>(Ct value) | HPV<br>(Ct value) |
| 1 | POS (35.9) | POS 21.5 | POS (32.3) | POS 23.2 | POS (34.7) | POS 22.8 | POS (30.9) | POS 22.8 | POS (33.3) | POS 20.6 |
| 2 | POS (32.4) | POS 25.6 | POS (31.9) | POS 28.7 | POS (32.3 ) | POS 26.9 | POS (30.6) | POS 26.8 | POS ( 32.4) | POS 23.1 |
| 3 | POS (32.3) | POS 29.1 | POS (31.1) | POS 30.4 | POS (33.9) | POS 30.8 | POS (30.5) | POS 29.0 | POS (31.8) | POS 27.6 |
| 4 | POS (31.7) | POS 31.7 | POS (31.4) | POS 34.9 | POS (33.4) | POS 34.8 | POS (30.9) | POS 32.3 | POS (31.3) | POS 31.0 |
| 5 | POS (32.3) | ND | POS (32.4) | ND | POS (32.9) | POS 38.9 | POS (31.1) | ND | POS (31.2) | POS 38.7 |
| 6 | POS (32.5) | ND | POS (31.3) | ND | POS (38.8) | ND | POS (31.7) | ND | POS (31.4) | ND |
| 7 | POS (35.9) | ND | POS (31.7) | ND | POS (35.3) | ND | POS (31.6) | ND | POS (31.3) | ND |
| HPV 18 |  |  |  |  |  |  |  |  |  |  |
| 1 | POS (25.9) | POS 22.9 | POS (26.1) | POS 23.7 | POS (26.4) | POS 25.8 | POS (28.7) | POS 26.6 | POS (25.9) | POS 23.3 |
| 2 | POS (29.5) | POS 27.0 | POS (28.5) | POS 27.2 | POS (29.0 ) | POS 29.4 | POS (30.4) | POS 28.7 | POS (28.9) | POS 27.3 |
| 3 | POS (30.1) | POS 31.0 | POS (30.1) | POS 31.1 | POS (29.3) | POS 31.5 | POS (30.1) | POS 29.6 | POS (30.1) | POS 31.7 |
| 4 | POS (29.3) | POS 33.9 | POS (30.4) | POS 38.4 | POS (30.1) | POS 38.4 | POS (29.5) | POS 33.0 | POS (31.6) | POS 35.4 |
| 5 | POS (30.3) | POS 39.7 | POS (29.2) | ND | POS (30.5) | ND | POS (30.5) | POS 36.0 | POS (30.5) | POS 38.9 |
| 6 | POS (29.8) | ND | POS (29.9) | ND | POS (29.6) | ND | POS (30.4) | ND | POS (30.3) | ND |
| 7 | POS (29.3) | ND | POS (29.7) | ND | POS (31.3) | ND | POS (31.2) | ND | POS (29.8) | ND |
| HPV 31 |  |  |  |  |  |  |  |  |  |  |
| 1 | POS (36.7) | POS 30.1 | POS (26.3) | POS 20.4 | POS (28.9) | POS 25.6 | POS (27.3) | POS 24.8 | POS (26.4) | POS 19.9 |
| 2 | POS (32.5) | POS 31.3 | POS (29.2 ) | POS 24.4 | POS (31.4) | POS ND | POS ( 29.9) | POS 28.3 | POS (30.1 ) | POS 24.8 |
| 3 | POS (32.5) | POS 33.8 | POS (29.5) | POS 26.8 | POS (30.2) | POS 31.8 | POS (32.5) | POS 37.2 | POS (30.5) | POS 29.0 |
| 4 | POS (32.4) | ND | POS (30.5) | POS 33.4 | POS (30.8) | ND | POS (30.2) | ND | POS (31.0) | POS 32.0 |
| 5 | POS (29.8) | POS 32.3 | POS (29.2) | ND | POS (30.4) | ND | POS (31.5) | ND | POS (31.3) | POS 36.1 |
| 6 | POS (30.1) | ND | POS (29.2) | ND | POS (28.1) | ND | POS (32.4) | ND | POS (30.8) | ND |
| 7 | POS (29.8) | ND | POS (29.9) | ND | POS (28.6) | ND | POS (31.3) | ND | POS (29.1) | ND |

SAC = sample adequacy control, Ct = Cycle threshold or point of DNA detection, ND= Not Detected

## References

1. Bray F, Ferlay J, Soerjomataram I, Siegel RL, Torre LA, Jemal A. Global cancer statistics 2018: GLOBOCAN estimates of incidence and mortality worldwide for 36 cancers in 185 countries. CA: a cancer journal for clinicians 2018; 68(6): 394–424.

2. Catarino R, Petignat P, Dongui G, Vassilakos P. Cervical cancer screening in developing countries at a crossroad: Emerging technologies and policy choices. World journal of clinical oncology 2015; 6(6): 281–90.

3. Dillner J, Rebolj M, Birembaut P, et al. Long term predictive values of cytology and human papillomavirus testing in cervical cancer screening: joint European cohort study. BMJ (Clinical research ed) 2008; 337: a1754.

4. Sankaranarayanan R, Nene BM, Shastri SS, et al. HPV screening for cervical cancer in rural India. N Engl J Med 2009; 360(14): 1385–94.

5. WHO. WHO guidelines for screening and treatment of precancerous lesions for cervical cancer prevention. South Africa: World Health Organization; 2013.

6. Einstein MH, Smith KM, Davis TE, et al. Clinical evaluation of the cartridge-based GeneXpert human papillomavirus assay in women referred for colposcopy. Journal of clinical microbiology 2014; 52(6): 2089–95.

7. Toliman PJ, Kaldor JM, Badman SG, Phillips S, Tan G, Brotherton JML, et al. Evaluation of self-collected vaginal specimens for the detection of high-risk human papillomavirus infection and the prediction of high-grade cervical intraepithelial lesions in a high-burden, low-resource setting. Clinical microbiology and infection: the official publication of the European Society of Clinical Microbiology and Infectious Diseases. 2019;25(4):496–503.

8. Arbyn M, Smith SB, Temin S, Sultana F, Castle P. Detecting cervical precancer and reaching underscreened women by using HPV testing on self samples: updated meta-analyses. BMJ (Clinical research ed) 2018; 363: k4823.

9. Cepheid - Xpert HPV package insert 301-2585, Rev B March 2014. Pp 4.

10. Hologic material data safety sheet. ThinPrep® PreservCyt Solution CPH MSDS NA 909897. Version #: 003. Issue date: 09-August-2013

11. Schrader, C, Schielke, A, Ellerbroek, L and Johne, R (2012), PCR inhibitors – occurrence, properties and removal. J Appl Microbiol, 113: 1014–1026.

12. Castle PE, Solomon D, Hildesheim A, Herrero R, Bratti MC, Sherman ME et al. 2003. Stability of archived liquid-based cervical cytologic specimens. Cancer 99:89–96

13. Negri G, Rigo B, Vittadello F, Egarter-Vigl E, Mian C. 2004. Human papillomavirus typing with hybrid capture II on archived liquid-based cytologic specimens: is HPV typing always reproducible? Am. J. Clin. Pathol. 122:90–93.

14. Castle PE, Hildesheim A, Schiffman M, Gaydos CA, Cullen A, Herrero R, et al. 2003. Stability of archived liquid-based cytologic specimens. Cancer 99:320–322.

15. Shipping ThinPrep® Solutions and Samples within CONUS 51-500 Vials – Limited Quantity Shipments. https://phc.amedd.army.mil/PHC%20Resource%20Library/ShippingThinPrep51-500Vials_FS_37-039-0518

